# Evolution and genetics of precocious burrowing behavior in *Peromyscus* mice

**DOI:** 10.1101/150243

**Authors:** Hillery C. Metz, Nicole L. Bedford, Linda Pan, Hopi E. Hoekstra

## Abstract

A central challenge in biology is to understand how innate behaviors evolve between closely related species. One way to elucidate how differences arise is to compare the development of behavior in species with distinct adult traits. Here, we report that *Peromyscus polionotus* is strikingly precocious with regard to burrowing behavior, but not other behaviors, compared to its sister species *P. maniculatus*. In *P. polionotus*, burrows were excavated as early as 17 days of age, while *P. maniculatus* did not build burrows until 10 days later. Moreover, the well-known differences in burrow architecture between adults of these species—*P. polionotus* adults excavate long burrows with an escape tunnel, while *P. maniculatus* dig short, single-tunnel burrows—were intact in juvenile burrowers. To test whether this juvenile behavior is influenced by early-life environment, pups of both species were reciprocally cross-fostered. Fostering did not alter the characteristic burrowing behavior of either species, suggesting these differences are genetic. In backcross F_2_ hybrids, we show that precocious burrowing and adult tunnel length are genetically correlated, and that a single *P. polionotus* allele in a genomic region linked to adult tunnel length is predictive of precocious burrow construction. The co-inheritance of developmental and adult traits indicates the same genetic region—either a single gene with pleiotropic effects, or closely linked genes— acts on distinct aspects of the same behavior across life stages. Such genetic variants likely affect behavioral drive (i.e. motivation) to burrow, and thereby affect both the development and adult expression of burrowing behavior.

**Highlights:**

- Juvenile *P. polionotus* construct burrows precociously compared to its sister species *P. maniculatus*
- Cross-fostering does not alter species-specific burrowing behavior
- A QTL linked to adult tunnel length predicts developmental onset of burrow construction in hybrids
- Pleiotropic genetic variant(s) may affect behavioral drive across life stages

## eTOC

Metz et al. find that oldfield mice, a species that digs long, complex burrows, also digs burrows earlier in development compared to its sister species. In F_2_ hybrids, precocious burrowing is co-inherited with long adult tunnels, and an allele linked to adult tunnel length also predicts timing of first burrow construction, suggesting that a single genetic region controls different aspects of the same behavior across distinct life stages.

## Results

### *P. polionotus* construct burrows earlier in life than *P. maniculatus*

To examine the developmental onset of burrow construction in *Peromyscus* mice, we assayed burrowing behavior in juveniles starting at 17 days of age (these mice are typically weaned at postnatal day [P] 24). We found striking interspecific differences in both the timing and progression of burrow construction (Figure 1; Table S1). Notably, *P. polionotus* were precocious diggers, constructing complete burrows—defined as excavations with at least two components: an entrance tunnel plus a nest chamber—on average 10 days earlier than *P. maniculatus*. The first appearance of a complete burrow was at P17 in *P. polionotus* (1 of 5 mice; Figure 1b), but not until P27 in *P. maniculatus* (3 of 14 mice; Figure 1b), a considerable difference in developmental stage (see Figure S1 for timeline of development). Moreover, *P. polionotus* burrowed at adult-like frequencies from P19 onward, a developmental benchmark *P. maniculatus* did not reach until P27 (Figure 1b; Table S1).

**Figure 1.**
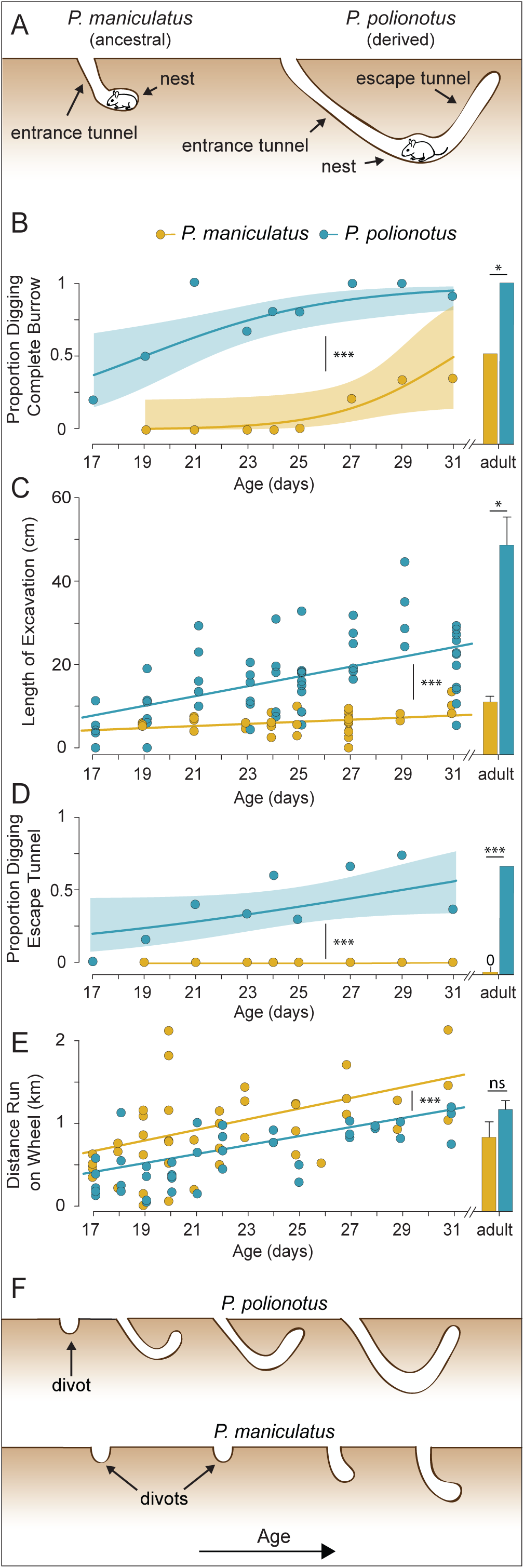
The ontogeny of burrow construction in sister species *P. maniculatus* (yellow) and *P. polionotus* (blue). All animals were naïve and tested only once. (**A**) The ancestral burrow architecture, built by *P. maniculatus*, is short (<15cm) and simple. In contrast, adult *P. polionotus* dig stereotyped burrows with a long entrance tunnel, nest chamber and escape tunnel (total excavation length ~50 cm). (**B**) Proportion of tested mice constructing a complete burrow (i.e., entrance tunnel and nest chamber). Curves and shaded areas represent binary generalized linear smoothers with 95% confidence intervals. Species differences evaluated by Fisher’s exact test (juveniles: see text for details, adults: *P. maniculatus* n=17; *P. polionotus* n=9). (**C**) Length of total excavation. Juvenile differences evaluated by ANCOVA (see text for details). Adult differences between species evaluated by *t*-test (*P. maniculatus* n=17; *P. polionotus* n=9). Error bars at +/- 1 SE of the mean. (**D**) Proportion of tested mice constructing an escape tunnel. Statistical tests as in (B). (**E**) Distance run on a wheel during a 90-minute trial by *P. maniculatus* and *P. polionotus* juveniles (P17-P31, see text for details) and adults (>P60, *P. maniculatus* n=10; *P. polionotus* n=10). Statistical tests as in (C). (**F**) Cartoon depiction of data shown in panels B-D highlighting variation in burrow shape over development. Significance levels: p ≥ 0.05 = ns (not significant), p ≤ 0.05 = *, p ≤ 0.01 = **, p ≤ 0.001 = ***.

Whereas tunnel length increased with age in both species, reflecting a progression in burrowing ability with growth and development (Figure 1c; ANCOVA, p < 0.0001), tunnel length varied considerably between species. *P. polionotus* consistently produced significantly longer burrows than *P. maniculatus* (Figure 1c; ANCOVA, p < 0.0001), consistent with the known differences in adult tunnel length [3-5]. Furthermore, the rate of increase in tunnel length across ontogeny was significantly greater for *P. polionotus* (Figure 1c; ANCOVA, age x species interaction, p = 0.030). Thus, both the expression of adult-like burrowing frequency and an increase in excavation length develops more rapidly in *P. polionotus* than in *P. maniculatus*.

In trials when mice did not construct full burrows, individuals of both species usually excavated shallow cup-shaped cavities (divots) instead. Only three of 97 mice (two P17 *P. polionotus* and one P27 *P. maniculatus*) failed to leave any signs of digging activity. These data suggest that the motor patterns for digging were partly, if not completely, developed in both species by at least P17.

### Juveniles construct burrows with miniaturized adult architecture

Juveniles from both species produced burrows with architecture typical of adults of their respective species. Starting at P19, nearly the earliest age tested, the burrows of *P. polionotus* included escape tunnels at a frequency not significantly different from conspecific adults (Figure 1d; Fisher’s exact test, p = 0.523). Likewise, *P. maniculatus* juvenile burrows invariably featured only a single tunnel leading to the nest chamber, always lacking an escape tunnel (Figure 1d). Although complete with regard to architectural components, juvenile excavations were significantly shorter than those of adults (Figure 1c; *t*-tests, p < 0.0001 for both species), thus representing miniature versions of adult burrows.

### Precociousness is specific to burrowing behavior

To evaluate whether precocious burrow construction in *P. polionotus* might be due to advantages in physical rather than behavioral development (e.g. [6]), we examined general measures of morphological and motor development in both species. Two lines of evidence refute this hypothesis. First, *P. polionotus* did not perform better in a second motor activity task: *P. polionotus* juveniles travelled less distance in a 90- minute wheel-running assay than *P. maniculatus*. While total distance run increased with age at a comparable rate in both species (Figure 1e; age × species interaction term, p = 0.5993), *P. maniculatus* ran significantly greater distances than age-matched *P. polionotus* (ANCOVA, p < 0.001). Second, *P. polionotus* are smaller than *P. maniculatus* in both body mass (ANCOVA, p < 0.0001) and hindfoot length (ANCOVA, p < 0.0001) across development (Figure S1). Likewise, we did not observe heterochrony favoring *P. polionotus* with respect to additional developmental milestones, as *P. maniculatus* reached them earlier in life (Figure S1). Thus, precocious burrowing in *P. polionotus* juveniles reflects a behavioral difference, likely specific to burrowing, not an advantage in overall activity level, motor ability, or morphological development.

### Species-specific burrowing behavior unaltered by interspecific cross-fostering

To disentangle the effects of genetics from environment, pups were reciprocally cross-fostered between the two sister species (Figure 2a). We reasoned that any effects on burrowing behavior resulting from parental environment were likely to be greatest during post-natal development.

**Figure 2.**
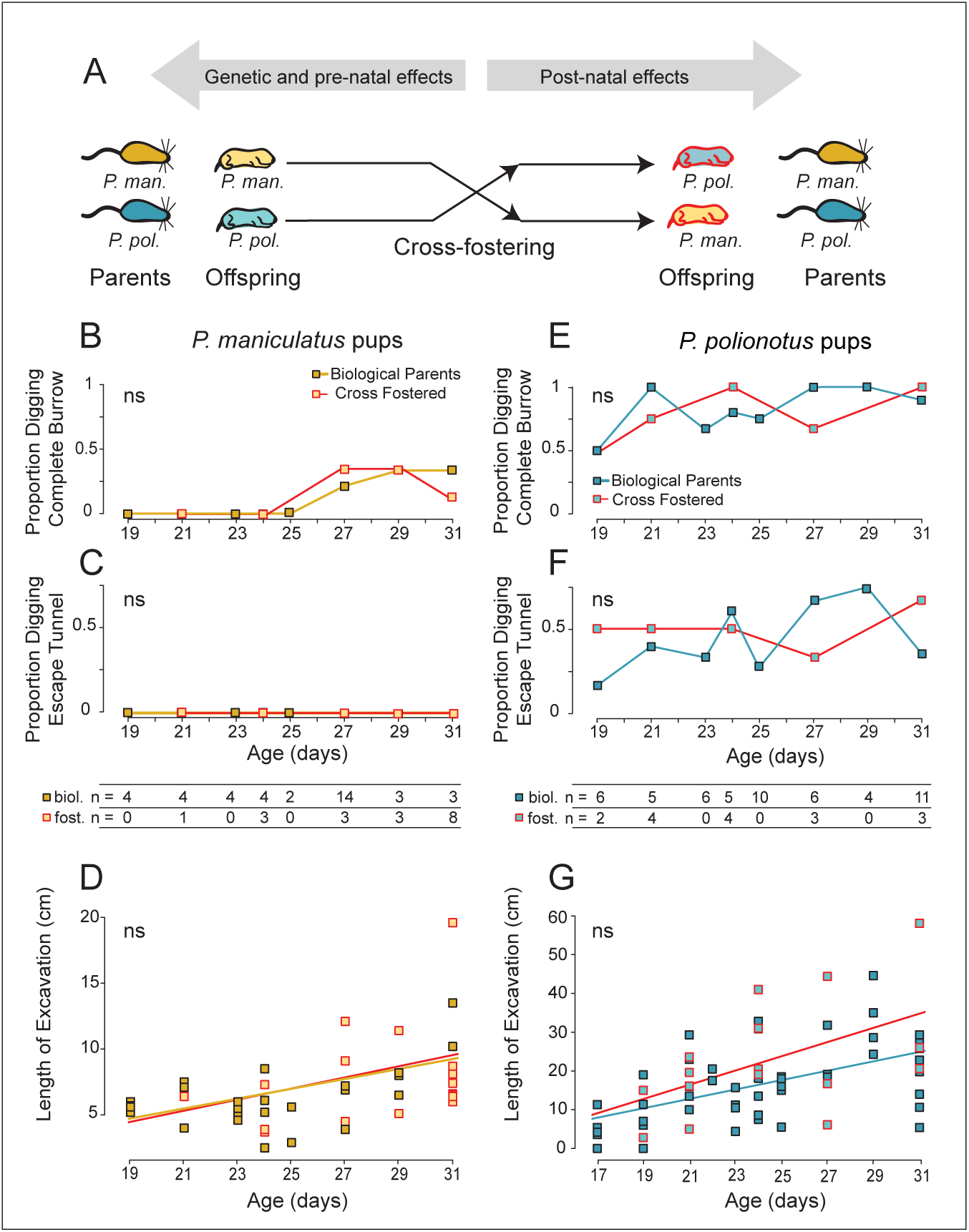
Reciprocal interspecific cross-fostering between *P. maniculatus* (yellow) and *P. polionotus* (blue). (**A**) Schematic of cross-fostering design with cross-fostered pups highlighted in red. (**B**,**E**) Proportion of mice constructing complete burrows, (**C**,**F**) proportion of mice building an escape tunnel, and (**D**,**G**) length of excavations. Sample sizes for each age and foster group are shown. For B, C, E and F, differences between foster treatments were evaluated by Fisher’s exact test; for D and G, by ANCOVA (see text for details). Significance levels: p ≥ 0.05 = ns (not significant).

In *P. maniculatus*, the developmental onset of burrow building did not differ between cross-fostered and non-fostered animals. Prior to P27, *P. maniculatus* juveniles did not build complete burrows regardless of foster treatment (Figure 2b). Following the onset of burrowing, fostered animals constructed burrows no more frequently than pups reared by their biological parents (Figure 2b; Fisher’s exact test, one-tailed, p = 0.57). Cross-fostered *P. maniculatus* also did not build escape tunnels (Figure 2c), and the excavations of cross-fostered animals closely matched those of mice raised by their biological parents with regard to length (Figure 2d; ANCOVA, p = 0.63).

Likewise, *P. polionotus* raised by heterospecific parents began burrowing at the earliest age tested (P19; Figure 2e), and from P21 onward, nearly all cross-fostered *P. polionotus* excavated burrows (12 of 14 mice; Figure 2e). Burrow structure also did not change with cross-fostering treatment. Cross-fostered *P. polionotus* dug escape tunnels as early in ontogeny (from P19), and as frequently (50%, 8 of 16 mice), as non-fostered juveniles (41%, 22 of 53 mice; Figure 2f, Fisher’s exact test, p = 0.34) and as frequently as conspecific adults (67%, 6 of 9 mice; Fisher’s exact test, p = 0.58). Finally, excavation lengths did not differ between cross-fostered and non-fostered animals (Figure 2g; ANCOVA, p = 0.06). In summary, we found no differences in burrowing behavior following cross-fostering, consistent with there being a strong genetic component to the development of burrowing behavior.

### Ontogeny of burrow construction is *P. polionotus*-dominant

We next tested the hypothesis that differences in the developmental onset of burrowing in juveniles share a common genetic basis with the well-characterized differences in adult burrow architecture [3-5] using a *P. polionotus* x *P. maniculatus* experimental cross (Figure 3a).

**Figure 3.**
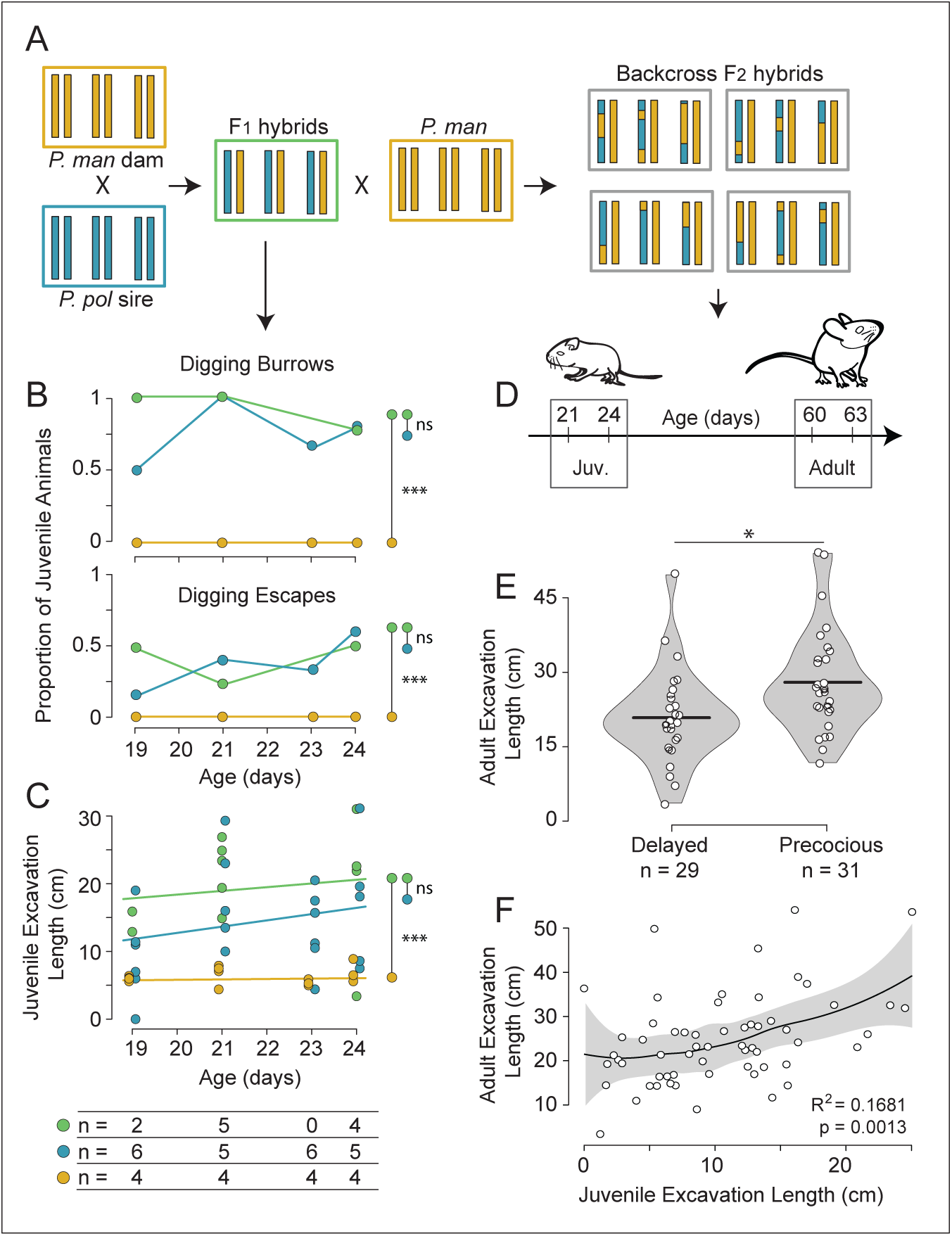
Genetic dissection of precocious burrowing in *P. polionotus* X *P. maniculatus* hybrids. (**A**) Schematic of breeding design showing *P. maniculatus* (yellow), *P. polionotus* (blue), first-generation F_1_ hybrids (green) and second-generation F_2_ hybrids (grey). (**B**) Proportion of juvenile animals digging complete burrows (upper panel) and escape tunnels (lower panel); groups compared using Fisher’s exact tests (see text for details). (**C**) Length of excavations in F_1_ hybrids compared to *P. maniculatus* and *P. polionotus*; species differences were evaluated by ANCOVA (see text for details). Sample sizes for each group are shown below. (**D**) Timeline of the four behavioral assays completed for each F_2_ hybrid. (**E**) Average adult excavation length of backcross F_2_ hybrids that, as juvenile burrowers, were either delayed (i.e., no burrows dug at P21 or P24) or precocious (i.e., at least one complete burrow dug at P21 or P24). Black lines indicate means for each group, which were compared by Welch’s two-sample *t*-test. (**F**) Least squares regression between juvenile and adult excavation lengths in backcross F_2_ hybrids (*R*^2^ = 0.1681, p = 0.0013). Shaded area is a Loess smoother (locally weighted smoother). Significance levels: p ≥ 0.05 = ns (not significant), p ≤ 0.05 = *, p ≤ 0.01 = **, p ≤ 0.001 = ***.

The development of burrowing behavior in first generation (F_1_) hybrids closely matches *P. polionotus* in each parameter examined, including the proportion of mice constructing burrows (Figure 3b; Fisher’s exact test, p = 0.378), the proportion of mice constructing escape tunnels (Figure 3b; Fisher’s exact test, p = 1.00), and the length of excavations (Figure 3c; ANCOVA, p = 0.086). Moreover, F_1_ hybrids differ significantly from *P. maniculatus* in all of these measures of burrowing behavior: proportion of mice constructing burrows (Figure 3b; Fisher’s exact test, p < 0.0001), proportion of mice constructing escape tunnels (Figure 3b; Fisher’s exact test, p = 0.019), and length of excavations (Figure 3c; ANCOVA, p < 0.0001). This inheritance pattern indicates that the genetic underpinnings of precocious burrowing, a developmental trait, are *P. polionotus*-dominant, consistent with the pattern of inheritance observed for adult burrowing behavior (F_1_ hybrid adults build *P. polionotus*-like burrows with regard to both length and shape [3,5]).

### Developmental and adult traits share a common genetic basis

To test if developmental traits (namely, precocious burrow construction) and adult traits (long entrance tunnels, presence of an escape tunnel) are genetically linked, we generated 60 backcross F_2_ hybrids. If traits have an independent genetic basis, they are expected to become uncoupled in the F_2_ generation. We assessed burrowing performance for each F_2_ hybrid at four time points: two juvenile (P21 and P24) and two adult trials (P61 and P64) (Figure 3d). Half of the F_2_ hybrids (31 of 60) dug at least one juvenile burrow (at the P21 or P24 time point) and thus were scored as precocious burrowers, while the remaining half (29 of 60) completed no juvenile burrows and were scored as delayed burrowers. Consistent with there being a shared genetic basis for burrowing traits across life stages, we found that precocious burrowers went on to dig significantly longer burrows as adults (Figure 3e; *t*-test, p = 0.024). Furthermore, juvenile excavation length was significantly associated with adult tunnel length (Figure 3f; least-squares linear regression, R^2^ = 0.1681, p = 0.001). Neither cross direction (i.e. whether the F_1_ parent was the sire or dam) nor sex of the F_2_ had an effect on juvenile or adult burrow length (*t*-tests, cross direction, juveniles: p = 0.371; adults: p = 0.673; sex, juveniles: p = 0.682; adults: p = 0.431). Together, these data suggest that for burrowing, juvenile (precocious onset of burrowing) and adult (tunnel length) behavior share some pleiotropic genetic basis, are influenced by closely linked genes, or both.

To determine which regions of the genome affect both juvenile and adult behaviors, we genotyped backcross F_2_ mice at four unlinked markers, each representing a genetic locus previously linked to differences in adult burrow architecture [5]. We found that inheritance of a *P. polionotus* allele was predictive of juvenile phenotype at one of the four markers (Figure 4). Specifically, a marker on linkage group 2 was associated with earlier onset of burrowing (Figure 4a; Fisher’s exact test, one-tailed, p = 0.044), juvenile excavation length (Figure 4b; *t*-test, p = 0.014), and, as expected, adult excavation length (Figure 4c; *t*-test p = 0.033). For each of the other markers examined, no significant relationships between genotype and phenotype were detected (Figure S2; Fisher’s exact tests and *t*-tests, p > 0.05), possibly due, in part, to the limited number of F_2_ hybrids examined. These data suggest that a gene, or closely linked genes, on linkage group 2 affects variation in burrowing behavior at different life stages.

**Figure 4.**
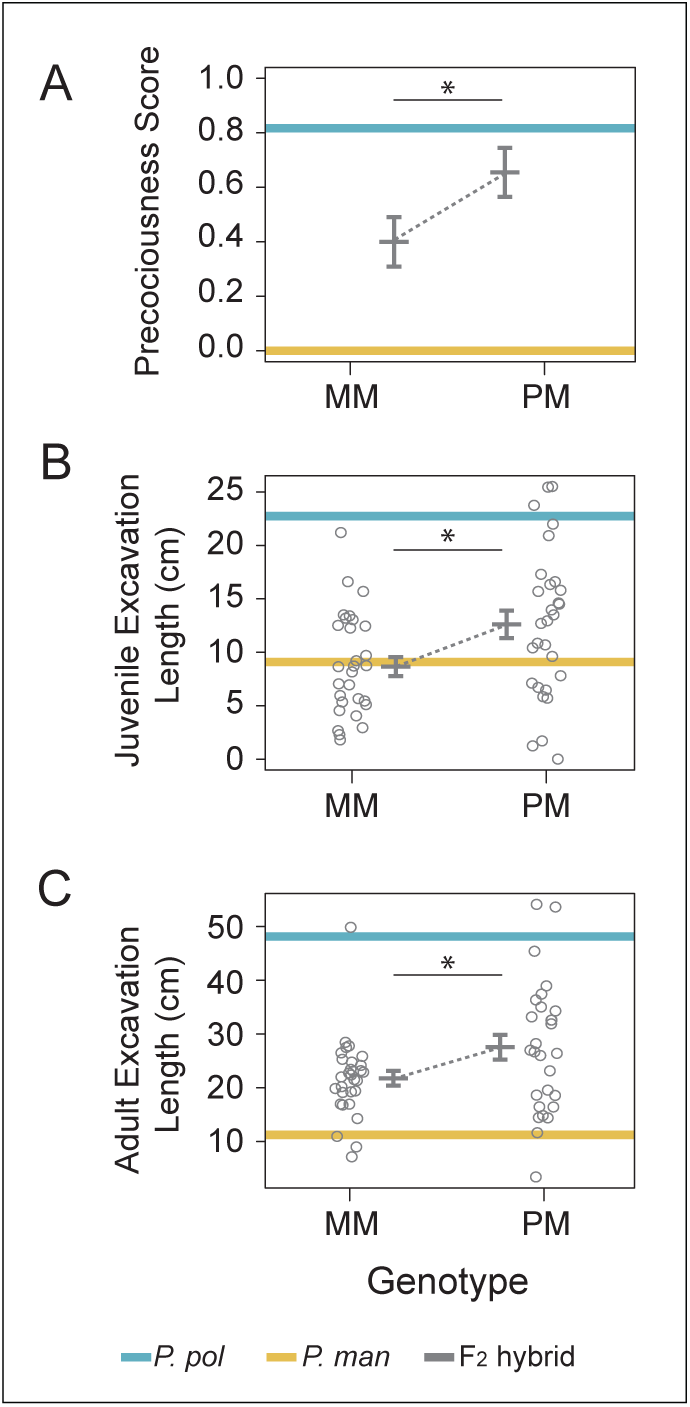
Effect of a single *P. polionotus* allele on juvenile burrowing behavior in F_2_ hybrids at a QTL previously linked to adult tunnel length [17]. Three traits are shown: (**A**) precociousness score across two juvenile trials (1 = mouse dug at least one discrete burrow at P21 and/or P24 behavior trials; 0 = mouse dug no burrow at P21 or P24); (**B**) average juvenile excavation length; and (**C**) average adult excavation length. Genotypes for 60 backcross F_2_ hybrids are either MM (homozygous for the *P. maniculatus allele*) or PM (heterozygous). For each genotype, trait means are plotted with error bars at *+/-* 1 SE of the mean. The mean trait values for each parental species are plotted as horizontal bars: *P. polionotus* (blue) and *P. maniculatus* (yellow). Parental species trait values are based on one trial per individual, aged P21-P24 (juveniles, *P. maniculatus* n=16; *P. polionotus* n=22), or >P60 (adults, *P. maniculatus* n=17; *P. polionotus* n=9). F_2_ hybrid trait values are the average of two juvenile (A,B) or two adult (C) trials. Significance levels, determined by Fisher’s exact test (A) or *t*-test (B,C), are: p ≤ 0.05 = *.

## Discussion

> *Huxley likes to speak of ‘the three major problems of biology’: that of causation, that of survival value and that of evolution—to which I should like to add a fourth, that of ontogeny.*
>
> *—Tinberg en (1963)*

Striking behavioral differences between closely-related species can be a powerful resource for understanding the evolution of behavior and its mechanistic underpinnings—both major goals of biology. Behaviors are among the most complex phenotypes, and to successfully tease apart how species-specific differences evolve requires an integrative approach, as championed by Tinbergen [7]. More specifically, Tinbergen’s 1963 landmark paper advocates for the addition of ontogeny to Huxley’s existing framework for behavioral research [8].

Ontogeny, the study of how behavior changes across the life of an individual, can provide understanding that is not discernible using other approaches; for example, it can uncover unexpected ancestral state reconstructions and generate novel hypotheses (e.g. [9-12]), or expose underlying proximate mechanisms driving changes in behavior (e.g. [13-16]). In short, ontogeny informs and edifies each of Tinbergen’s four questions and can provide novel insights into how behavior evolves.

Here, we focused on the ontogeny of burrow construction, an ecologically important behavior that varies dramatically between closely related species of North American *Peromyscus* rodents. Most species in this genus build small (<20cm), simple burrows as adults, but one species, *P. polionotus,* has recently evolved a stereotyped burrowing behavior that results in a considerably longer burrow (>100cm in the wild) comprised of an elongated entrance tunnel, a nest chamber, and a secondary tunnel that extends upward from the nest toward the soil surface. This second tunnel does not penetrate the soil surface except during emergency evacuation, and thus is often referred to as an escape tunnel ([3-5, 17-20]; Figure 1a). The burrows of *P. polionotus* have inspired studies of phylogenetic history [4], genetic mechanisms of behavior [3,5], and speculations of adaptive function—namely that *P. polionotus* burrows may provide refuge from the elevated rates of predation that occur in open, exposed habitats (e.g. [21,22]). However, the ontogeny of the behavior—the last of Tinbergen’s four questions—remained unexamined until now.

We report on how the final product of digging behavior—the extended phenotype [23], or burrow—originates and progresses during the post-natal development of two sister species of *Peromyscus* with dramatically different adult burrow architectures. We first find that *P. polionotus* are precocious with respect to burrow construction, building their first burrows 10 days earlier in development than *P. maniculatus*. This is surprising given that *P. maniculatus* is larger, tends to reach developmental milestones earlier, and outperforms age-matched *P. polionotus* in a wheel-running assay. These results suggest that *P. polionotus* has evolved a life history change—a precocious expression of behavior—that is likely specific to burrow construction.

We also examined the shape of burrows produced by juvenile *Peromyscus* mice. We found that each species’ characteristic burrow architecture is intact in juveniles. This result suggests that in pure species, the neurobiological control of each component of the complete burrow architecture (frequency of burrow construction, entrance tunnel, and escape tunnel) is expressed together throughout life. This result is especially surprising in light of previous work showing that the genetic control of adult burrow construction in *P. polionotus* is modular [5]. Although the shape of juvenile burrows is similar to adult burrows, they are smaller in overall size, likely due to the energetic cost of burrowing.

Using a cross-fostering experiment, we next tested if these juvenile burrowing traits were primarily learned postnatally or were driven by interspecific genetic differences. It is important to note, however, that our experiments cannot rule out prenatal maternal effects (e.g. [24]). We found that cross-fostering results do not differ if single or multiple pups are transferred to heterospecific parents, suggesting there is no measurable effect of sibling’s genotype on juvenile behavior. We report that all aspects of species-specific burrowing behavior are preserved in cross-fostered individuals of both species, demonstrating that juvenile expression of burrowing behavior is likely to have a strong genetic basis.

Finally, we examined the genetic underpinnings of behavioral ontogeny in hybrids of *P. polionotus* and *P. maniculatus* using a genetic cross. We found that a developmental trait (precocious onset of burrowing) and an adult trait (long tunnels characteristic of adult *P. polionotus* burrows) are genetically dominant and co-inherited, both at the level of phenotypic co-variation and with respect to a specific genetic marker. These data point to a shared—likely pleiotropic—genetic basis influencing behavior across life stages.

These results have implications for the evolution of burrowing behavior. First, pleiotropy (or tight linkage of multiple causal mutations) can facilitate or inhibit evolution. On one hand, pleiotropy can produce effects that are not directly selected for (and potentially even harmful), but that are nevertheless secondarily “dragged along” by evolution [25,26]. On the other hand, because changes in several traits are often involved during adaptation to a new environment [27-29], co-inheritance of groups of phenotypes (e.g. by pleiotropy or linkage) can expedite adaptation [30-32]. Indeed, a common experimental outcome is to map multiple traits to a shared genomic region [33-38], and this genetic architecture can affect how evolution proceeds.

Related, these findings make it difficult to identify the precise phenotypic targets of selection, if any. While variation in adult burrows can affect fitness [22,39], juvenile burrowing behavior may also be a target of selection. For example, natural selection for burrowing earlier in *P. polionotus* may reflect (i) its open habitat [17], which may expose young mice to predation and thus increase the survival value of burrowing, or (ii) a form of “play” during a critical period of motor development [40-42]. Our results, which implicate a broadly-acting pleiotropic genetic mechanism, highlight the challenge in identifying which specific trait or traits have been selected—in this case, precocious juvenile burrowing, long adult burrows, or both.

Finally, the shared genetic control of the timing of behavior onset and adult behavior also sheds light on the underlying neural mechanism. One parsimonious explanation for the co-inheritance of precocious burrowing with long entrance tunnels is that species-specific genetic differences produce heritable internal states that persist in individuals across life stages. Specifically, genetic variation shaping internal states may affect innate, species-specific behavioral drives or time budgets (i.e. apportionment of time spent performing different behaviors) such that *P. polionotus* more frequently engages in burrowing rather than alternative behaviors, compared to *P. maniculatus,* whose innate drives are tuned differently). Divergent neural tuning often has been linked to variation in neuromodulators or their receptors, rather than to variation in the underlying circuitry (e.g. [43-46]). Our results are thus consistent with a role for neuromodulators (and behavioral drives) in the evolution of burrowing in *Peromyscus* rodents, adding to the accumulating evidence that neuromodulatory systems are a frequent substrate for behavioral diversity and evolution [47].

## Experimental Procedures

### Animal care and breeding

We used captive strains of *Peromyscus* originally acquired from the *Peromyscus* Genetic Stock Center (University of South Carolina, Columbia SC, USA). We used only offspring of experienced parents (≥1 previous litter weaned) for experiments. Mice were housed in standard conditions. Due to genomic imprinting in these species, production of F_1_ hybrids was limited to crosses of *P. polionotus* sires to *P. maniculatus* dams [48,49]. Thus, this cross design excludes any *P. polionotus* maternal effects acting in favor of *P. polionotus*-like burrowing behavior. All procedures were approved by the Harvard University Institutional Animal Care and Use Committee.

### Burrowing behavior trials: parental species and F_1_ hybrids

We tested burrowing behavior in a total of 131 juvenile and 26 adult mice in in large, indoor sand-filled arenas as previously described [4,5], except duration was reduced from 48 hours to 14-17 hours for juveniles. Briefly, we released animals into 1.2 × 1.5 × 1.1 m enclosures filled with approximately 700 kg of moistened, hard-packed premium playground sand (Pharma-Serv Corp.), under otherwise normal housing conditions. We tested juveniles once, without previous experience, and thus our experiment targeted innate behavior and not learned ability. Each mouse was tested once, singly.

### Burrowing behavior trials: backcross F_2_ hybrids

We generated 60 backcross F_2_ hybrids by crossing F_1_ hybrids to *P. maniculatus* mates (Figure 3a). Both male and female F_1_ hybrids were backcrossed to *P. maniculatus* (reciprocal pairings): 22 animals were produced from an F_1_ dam and 38 from an F_1_ sire. We then characterized the juvenile and adult burrowing behavior of these backcross mice, collecting developmental and adult phenotypes in the same individuals: each F_2_ was tested four times in total, at juvenile ages 21 and 24 days, and adult ages 61 and 64 days. Enclosure area was reduced by half (i.e. to 0.6 × 1.5 × 1.1 m) for testing both juvenile and adult backcross individuals to accommodate the large number of animals being tested.

### Burrow Measurements

To quantify burrow construction, at the conclusion of each trial, we inspected enclosures for any excavations, which were qualitatively characterized as either burrows (comprised of ≥1 tunnel plus a nest area) or divots (broad cup-shaped vertical diggings <10 cm; see Figure 1f). Next, we injected burrows with polyurethane insulation foam (Hilti Corp., Schaan, Liechtenstein) as previously described [4,5] and measured the lengths of burrow components (entrance tunnel, nest chamber, and escape tunnel if present) from dried polyurethane casts. We measured the length of divots directly in the enclosures.

### Cross-fostering

Age-matched (≤ 48 hrs age difference) *P. maniculatus* (n=18) and *P. polionotus* (n=16) pups were traded between experienced (≥1 previous litter) heterospecific breeding pairs 24-48 hours after birth. Then, we measured the burrowing behavior of each resultant juvenile at a single time point (during 19-31 days).

### Wheel-running Behavioral Trials

To compare the ontogeny of a second motor behavior (and general activity level) between species, we performed a standardized wheel running assay [50]. We tested naïve, juvenile *P. maniculatus* (n=43) and *P. polionotus* (n=40) at P17 - P31. After 4 hours of home cage habituation to the wheel (Ware Manufacturing Inc., Phoenix, AZ), we recorded 90 minutes of wheel running activity with a CC-COM10W wireless bike computer (Cateye Co. Ltd., Osaka, Japan). We performed all tests between 16:00 and 22:00h during the dark cycle.

### Statistical Analyses

To disentangle effects of age on burrowing behavior, we employed several statistical tests. We first tested for effects of age and species on burrowing behavior as well as for effects of sex, postnatal litter size, enclosure, and foster status at the intraspecific level using ANCOVA. Because we did not detect statistical differences between treatments, singly cross-fostered individuals and litter-fostered animals were pooled (fostering details above). We used Fisher’s exact test to evaluate the frequencies of burrow and escape tunnel digging between different genetic groups. We used *t*-tests to compare means in F_2_ hybrids. To evaluate phenotypic correlations in our F_2_ cross, we used least squares linear regression. To detect associations between quantitative trait loci (QTLs) linked to adult behavior and precocious burrowing, we used Fisher’s exact test, and for continuous traits in juveniles (i.e. excavation length), we used *t*-tests. Two *P. polionotus* individuals that appeared in poor health (age >23 days) were excluded from all analyses. All statistical analyses were performed using the R language.

### Genotyping

We genotyped the backcross F_2_ population (n=60) at four loci corresponding to known QTL underlying adult burrowing behavior (identified in [5]) using species-specific restriction fragment length polymorphism (RFLP) differences. We verified that the selected RFLPs were fixed between species by sequencing PCR amplicons of 4 unrelated individuals of each species as well as the *P. maniculatus* and F_1_ parents of the cross (BigDye® Terminator v3.1 Cycle Sequencing Kit, Life Technologies). PCR products were digested with restriction enzymes (New England Biolabs, Ipswich MA), separated by gel electrophoresis, and genotypes were called based on the resultant banding pattern.

## Author Contributions

H.C.M., N.L.B. and H.E.H. conceived and designed experiments. H.C.M., N.L.B. and L.P. performed experiments, H.C.M. and N.L.B. analyzed the data. H.C.M. and H.E.H wrote the paper.

## Acknowledgements

We thank S. Scalia, J. Mason and the Hoekstra Lab for assistance with behavioral assays and animal husbandry; A. Bendesky, B. de Bivort, C. Dulac, J. Losos, B. Ölveczky, H. Fisher, B. Peterson, M. Munoz, and J. Weber for helpful discussions and comments on the manuscript; Harvard’s Office of Animal Resources, particularly J. Rocca for animal husbandry; and S. Worthington for statistical consultation. This research was funded by a Chapman Memorial Scholarship, a National Science Foundation Graduate Research Fellowship, a Doctoral Dissertation Improvement Grant (NSF 1209753), and an American Fellowship from the American Association of University Women to H.C.M.; a Natural Sciences and Engineering Research Council of Canada Graduate Fellowship to N.L.B.; Harvard College Research Program award and a Harvard Museum of Comparative Zoology Grant for Undergraduate Research to L.P., and a Beckman Young Investigator Award to H.E.H. H.E.H. is an Investigator in the Howard Hughes Medical Institute.

## Supplementary Figure Legends

**Figure S1.**
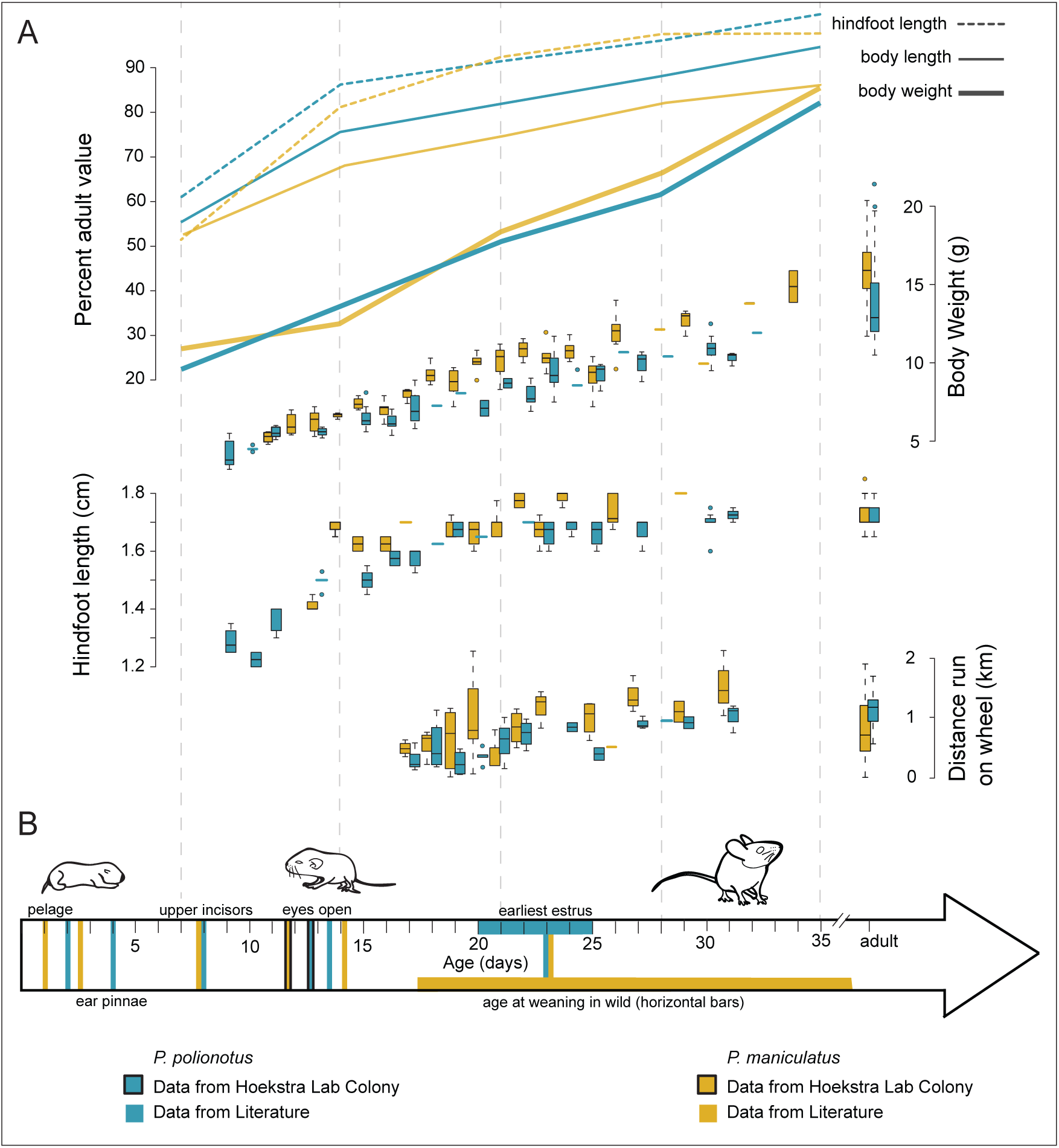
Comparison of morphological and motor development in *P. polionotus* (blue) and *P. maniculatus* (yellow) juveniles. (**A**) Morphological growth as a percentage of adult body weight, body length and hindfoot length (data adapted from[1]), as well as absolute body weight, absolute hindfoot length, and distance running on a wheel. (**B**) Ontogenetic trajectory of *P. polionotus* and *P. maniculatus* at key developmental events (data adapted from [1,2] as indicated).

**Figure S2.**
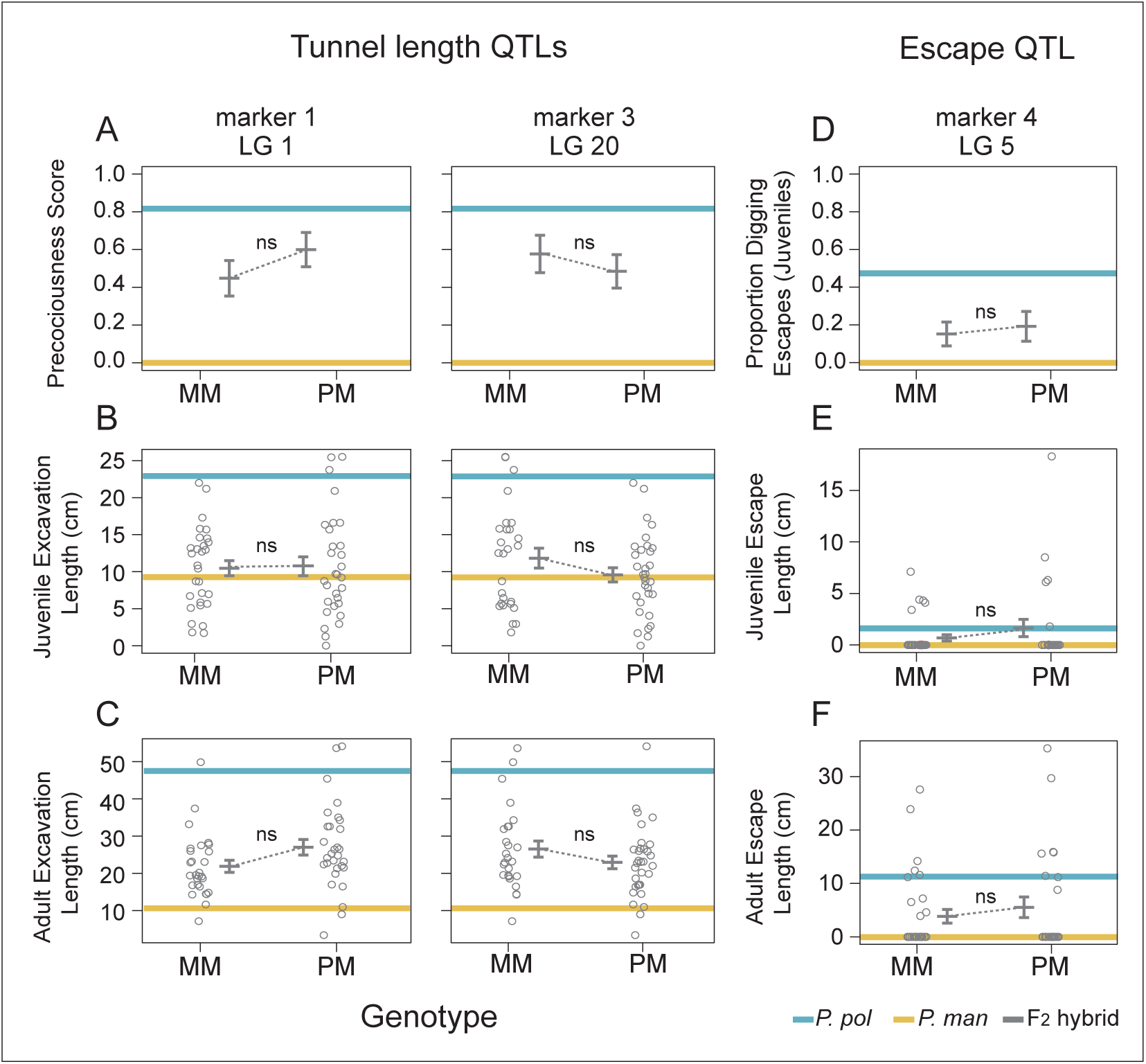
Effect of genotype on juvenile burrowing behavior in F_2_ hybrids at three additional QTLs previously linked to adult burrowing behavior [5]. Six total traits are shown. Two QTL are associated with adult burrow length (linkage groups 1 and 20): (**A**) precociousness score across two trials (1 = mouse dug at least one discrete burrow during P21 and P24 behavior trials; 0 = mouse dug no burrows at P21 or P24); (**B**) average juvenile excavation length; (**C**) average adult excavation length. One QTL (linkage group 5) was associated previously with escape tunnel construction in adults: (**D**) proportion of juveniles constructing escape tunnels; (**E**) juvenile escape tunnel length and (**F**) adult escape tunnel length. Genotypes for 60 backcross F_2_ hybrids are either MM (homozygous for the *P. maniculatus allele*) or PM (heterozygous). For each genotype, trait means are plotted with error bars at *+/-* 1 SE of the mean. The mean trait values for each parental species are plotted as horizontal bars: *P. polionotus* (blue) and *P. maniculatus* (yellow). Parental species trait values are based on one trial per individual, aged P21-P24 (juveniles, *P. maniculatus* n=16; *P. polionotus* n=22), or >P60 (adults, *P. maniculatus* n=17; *P. polionotus* n=9). F_2_ hybrid trait values are the average of two juvenile (A, B, D, E) or two adult (C, F) trials. Significance levels, determined by Fisher’s exact test or *t*-test, are: p ≥ 0.05 = ns (not significant).

**Supplemental Table 1.** Proportion of mice digging complete burrows (i.e. entrance tunnel and nest chamber) at three stages: pre-weaning (ages 19-25 days), weaned juveniles (ages 27-31 days) and adults (60+ days of age).

|  | *P. pol*<br>27-31 days<br>(n=21) | *P. pol*<br>Adult<br>(n=9) | *P. man*<br>19-25 days<br>(n=19) | *P. man*<br>27-31 days<br>(n=20) | *P. man*<br>Adult<br>(n=17) |
| --- | --- | --- | --- | --- | --- |
| *P. pol*<br>19-25 days<br>(n = 32) | p = 0.0707 | p = 0.1645 | p < 0.0001<br>*** | p = 0.0006<br>*** | p = 0.1996 |
| *P. pol*<br>27-31 days<br>(n=21) |  | p = 1.00 | p < 0.0001<br>*** | p < 0.0001<br>*** | p = 0.0051<br>** |
| *P. pol*<br>Adult<br>(n=9) |  |  | p < 0.0001<br>*** | p = 0.0002<br>*** | p = 0.0233<br>* |
| *P. man*<br>19-25 days<br>(n=19) |  |  |  | p = 0.0471<br>* | p = 0.0003<br>*** |
| *P. man*<br>27-31 days<br>(n=20) |  |  |  |  | p = 0.1014 |

p-values for 2 x 2 Fisher’s exact tests. Significance levels: $p \leq 0.05 = *$ , $p \leq 0.01 = **$ , $p \leq 0.001 = ***$ .

**Supplemental Table 2.**
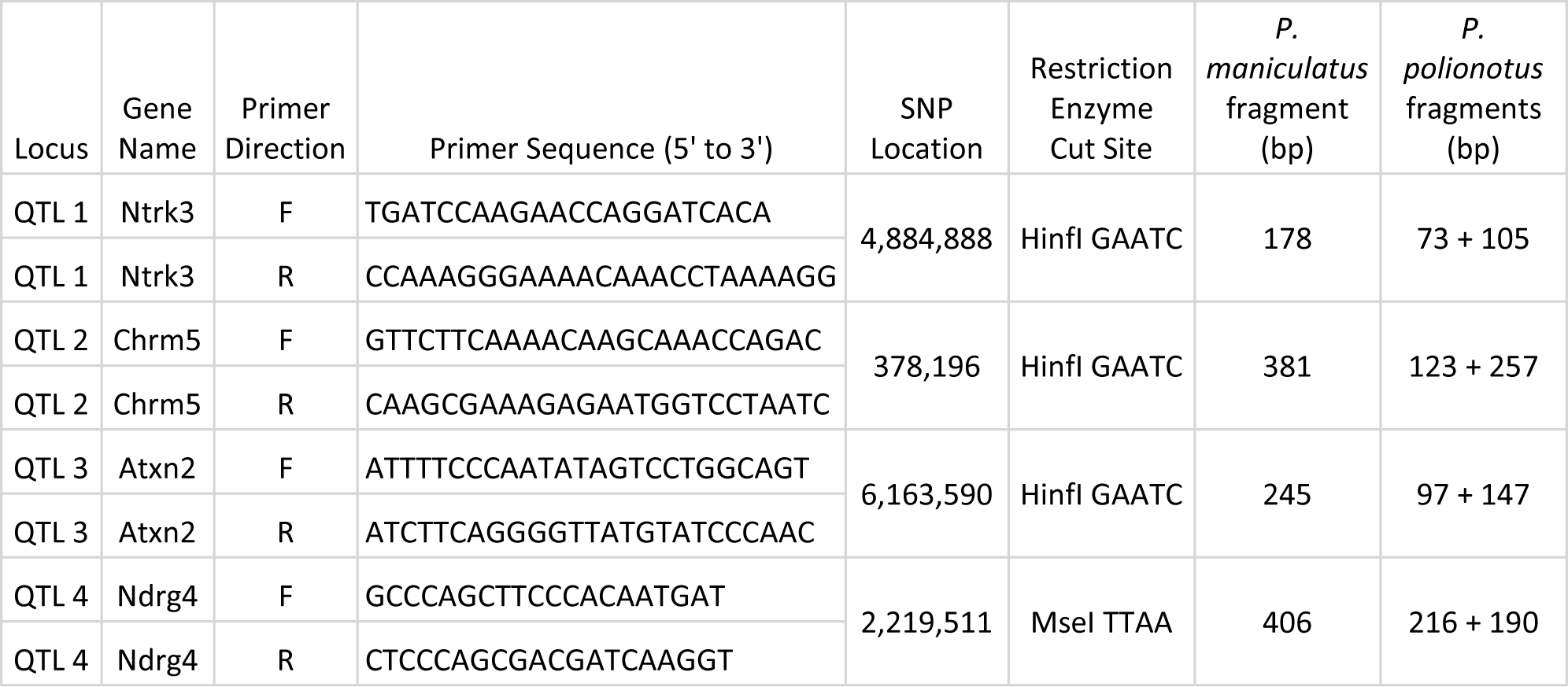
Primer sequences and restriction enzymes used for RFLP genotyping assays at four loci.

## Supplemental Materials and Methods

### Experimental Procedures

#### Animal care and breeding

We conducted experiments using captive *Peromyscus* strains kept under controlled laboratory conditions. Strains of both species were originally acquired from the *Peromyscus* Genetic Stock Center (University of South Carolina, Columbia SC, USA). These outbred lines, derived from wild-caught ancestors in 1948 (*P. maniculatus,* BW strain) and 1952 (*P. polionotus,* PO strain), have been laboratory-housed since capture, and have thus lived without access to soil for well over 100 generations. For each species, we formed breeding pairs using unrelated adults and checked daily for the presence of new pups. We used only offspring of experienced parents (≥1 previous litter weaned) for experiments. Mice were housed at 21.1°C on a 16:8 h light:dark cycle and were provided high-fat breeding diet and water *ad libitum*. We provided animals with access to cotton nesting material, corn cob bedding material, and 3- sided red-tinted plastic shelters. We weaned juveniles from their parents 24 days after birth, and weanlings were subsequently housed in littermate groups until completion of experiments. Due to genomic imprinting in these species, our cross design for production of F_1_ hybrids was limited to *P. polionotus* sires to *P. maniculatus* dams [48, 49]. Thus, this cross design excludes any *P. polionotus* maternal effects acting in favor of *P. polionotus*-like burrowing behavior. All procedures were approved by the Harvard University Institutional Animal Care and Use Committee.

#### Burrowing behavior trials: parental species and F_1_ hybrids

We tested the burrowing behavior in a total of 131 juvenile mice: 57 *P. maniculatus* (including 18 cross-fostered individuals) and 74 *P. polionotus* (16 cross-fostered) ranging in age from P17-P31, and 11 F_1_ hybrids ranging in age from P19-P24. For comparison, we also tested 17 adult *P. maniculatus* and 9 adult *P. polionotus* under similar conditions. We tested both males and females, but no sex-specific differences in burrowing behavior have been observed, here or previously [17].

We conducted all burrowing trials in large, indoor sand-filled arenas as previously described [4,5], except duration was reduced from 48 hours to 14-17 hours (one 8-hour dark cycle followed by 6-9 hours of light) for juveniles because the youngest mice could not endure an extended separated from their dam. Briefly, we released animals into large enclosures (1.2 × 1.5 × 1.1 m) filled with approximately 700 kg of moistened, hard-packed premium playground sand (Pharma-Serv Corp.). Mice were provided with nesting material, standard rodent food, and water during trials. Temperature and light:dark cycle were identical to housing conditions. For mice ≤ 24 days of age, we also provided Napa Nectar (Lenderking, MD, USA) and Nutrical (Tomlyn, SC, USA), as is standard practice for weanling-age animals. We tested juveniles once, without previous experience, and thus our experiment targeted innate behavior and not learned ability. Thus, each mouse was tested once, singly. We tested mice of both species at postnatal ages P19, P21, P23, P24, P25, P27, P29 and P31 days. Because of the species’ early onset of burrowing behavior, we tested additional *P. polionotus* individuals at P17, the earliest possible age to separate a juvenile from its mother. We tested F_1_ hybrids at P19, P21, and P24. Thus, we constructed a developmental time series for each species during key stages of motor and behavioral development.

#### Burrowing behavior trials: backcross F_2_ hybrids

We generated 60 backcross F_2_ hybrids by mating F_1_ hybrids to *P. maniculatus* mates (experimental design shown in Figure 3a). Both male and female F_1_ hybrids were backcrossed to *P. maniculatus* (reciprocal pairings): 22 animals were produced from F_1_ dam x *P. maniculatus* sire and 38 from *P. maniculatus* dam X F_1_ sire. We then characterized the juvenile and adult burrowing behavior of these backcross mice, thus measuring developmental and adult phenotypes in the same individuals. We tested juvenile mice at P21 and P24, when the greatest differences in juvenile burrowing behavior between *P. maniculatus* and *P. polionotus* are observed (Figure 1b). We tested these same mice again as adults (at P61 and P64) using the same assay. For all F_2_ backcross assays, we reduced the enclosure area by half (i.e. to 0.6 × 1.5 × 1.1 m) to accommodate the large number of animals being tested.

#### Burrow Measurements

To quantify burrow construction, at the conclusion of each trial, we inspected enclosures for any excavations, which were qualitatively characterized as either burrows (comprised of ≥1 tunnel plus a discernable nest area), or divots (broad cup-shaped vertical diggings <10 cm); see Figure 1f. Next, we injected burrows with polyurethane insulation foam (Hilti Corp., Schaan, Liechtenstein) as previously described [4,5], and measured the lengths of burrow components (entrance tunnel, nest chamber, and escape tunnel if present) from these dried polyurethane casts. We measured the length of divots directly in the sand enclosures.

#### Cross-fostering

To disentangle the effects of genetics from social environment during postnatal development, pups were reciprocally cross-fostered between the two species. We traded age-matched (≤ 48 hrs age difference) *P. maniculatus* (n = 18) and *P. polionotus* (n = 16) pups between experienced (≥1 previous litter) breeding pairs 24-48 hours after birth; the mice were otherwise kept under constant environmental and housing conditions. To test for effects of parents versus siblings on the behavior of the test animal(s), we used two fostering paradigms: pups were fostered as either singles (one pup traded between litters, such that the fostered pup had heterospecific siblings and heterospecific parents) or as litters (entire litters traded between breeding pairs, such that pups had heterospecific foster parents but conspecific siblings). Because the burrowing performance of both singly and group cross-fostered animals did not differ (ANCOVA; *P. polionotus*: age p = 0.010, foster treatment p = 0.880, age x treatment interaction p = 0.677; *P. maniculatus:* age p = 0.006, foster treatment p = 0.807, treatment x age interaction p = 0.853), we grouped these data together for subsequent analyses comparing fostered and non-fostered animals. Following weaning from their parents at P24, juveniles were housed with cage-mate siblings (biological or foster) until the beginning of behavioral trials.

#### Wheel-running Behavioral Trials

To compare the ontogeny of a second motor behavior (and general activity level) between species, we performed a standardized wheel running assay. We tested juvenile *P. maniculatus* (n = 43) and *P. polionotus* (n = 40) between P17 and P31. Both males and females were tested. We outfitted a 5-inch flying saucer exercise wheel (Ware Manufacturing Inc., Phoenix, AZ) with a CC-COM10W wireless bike computer (Cateye Co. Ltd., Osaka, Japan) to record total distance run over 90 minutes. To habituate the mouse to the novel object, we placed an exercise wheel in the home cage 4 hours prior to testing. We then placed the mouse in a new standard cage with a clean wheel and unlimited food and water. *Peromyscus* show strongly nocturnal patterns of wheel running [50], thus we performed all tests between 16:00 and 22:00h during the dark cycle.

#### Statistical Analyses

To disentangle effects of age on burrowing behavior, we employed several statistical tests. We first tested for effects of age and species on burrowing behavior as well as for effects of sex, postnatal litter size, enclosure, and foster status at the intraspecific level using ANCOVA. Because we did not detect statistical differences between treatments, we pooled singly cross-fostered individuals (having both heterospecific parents and siblings) and litter-fostered animals (heterospecific parents but conspecific siblings). We used Fisher’s exact test to evaluate the frequencies of burrow and escape tunnel digging between different genetic groups, and t-tests to compare means in F_2_ hybrids. To evaluate phenotypic correlations in our F_2_ cross, we used least squares linear regression. To detect associations between QTLs linked to adult behavior and precocious burrowing, we used both Fisher’s exact test, and t-tests for continuous traits in juveniles (i.e. excavation length). We excluded two *P. polionotus* individuals that appeared in poor health (age >23 days) from all analyses. We performed all statistical analyses using the R language.

#### Genotyping

We made use of restriction fragment length polymorphisms (RFLPs) to genotype the backcross F_2_ population (n = 60) at four loci corresponding to known QTL underlying adult burrowing behavior (QTL identified in [5]). We designed all four assays such that the *P. polionotus* allele contained a restriction enzyme cut site, whereas the *P. maniculatus* allele did not. We performed PCR with a Taq DNA Polymerase kit (Qiagen) and custom primers (Integrated DNA Technologies; Table S2). We verified that the selected RFLPs were fixed between species by Sanger sequencing (3730XL DNA Analyser, Applied Biosystems) of PCR amplicons in 4 unrelated individuals of each species as well as the *P. maniculatus* and F_1_ parents of the cross (BigDye® Terminator v3.1 Cycle Sequencing Kit, Life Technologies). Next, we digested PCR products with restriction enzymes (New England Biolabs, Ipswich MA) and separated the products by gel electrophoresis using a Quick-Load® 100bp DNA Ladder (New England Biolabs, Ipswich MA) as a size reference. We visualized digested fragments under UV light. All backcross animals inherit at least one *P. maniculatus* allele; therefore, we interpreted the presence of a second smaller fragment (of appropriate size) as evidence of a *P. polionotus* allele.

